# How to measure bacterial genome plasticity? A novel time-integrated index helps gather insights on pathogens

**DOI:** 10.1101/2024.01.22.576626

**Authors:** Greta Bellinzona, Gherard Batisti Biffignandi, Fausto Baldanti, Matteo Brilli, Davide Sassera, Stefano Gaiarsa

## Abstract

Genome plasticity can be defined as the capacity of a bacterial population to swiftly gain or lose genes. The time factor plays a fundamental role for the evolutionary success of microbes, particularly when considering pathogens and their tendency to gain antimicrobial resistance factors under the pressure of the extensive use of antibiotics. Multiple metrics have been proposed to provide insights into the gene content repertoire, yet they overlook the temporal component, which has a critical role in determining the adaptation and survival of a bacterial strain. In this study, we introduce a novel index that incorporates the time dimension to assess the rate at which bacteria exchange genes, thus fitting the definition of plasticity. Opposite to available indexes, our method also takes into account the possibility of contiguous genes being transferred together in one single event. We applied our novel index to measure plasticity in three widely studied bacterial species: *Klebsiella pneumoniae*, *Staphylococcus aureus*, and *Escherichia coli*. Our results highlight distinctive plasticity patterns in specific sequence types and clusters, suggesting a possible correlation between heightened genome plasticity and globally recognized high-risk clones. Our approach holds promise as an index for predicting the emergence of strains of potential clinical concern, possibly allowing for timely and more effective interventions.

**Impact statement:** How quickly bacterial populations can acquire new functions is the key to their evolutionary success. This speed, called genome plasticity, is particularly relevant for human pathogens, especially when considering the acquisition of antimicrobial resistance. Today, the availability of large numbers of genomes from public databases makes it possible to develop a way to measure plasticity. However, none is currently available, besides indexes of gene content variability, which do not take into account the rate at which such gene content changes. In this work, we developed a plasticity index, called Flux Of Gene Segments (FOGS), and we tested it on large datasets of bacterial pathogen genomes. Interestingly, the subpopulations of the selected species that showed higher FOGS correspond to globally emerging high-risk clones. Therefore, we suggest that our index might be used not only to detect but also to predict emerging strains of human health concern.

**Data summary:** The authors confirm that all supporting data, code and protocols have been provided within the article or through supplementary data files.

## Introduction

The evolutionary success of an organism hinges on its capacity to continuously adapt to changes encountered within the environment. Prokaryotes are notably characterized by the ability to acquire exogenous DNA through horizontal gene transfer (HGT) [1] and easily lose unnecessary genes [2], both representing powerful adaptation tools. The balance between gene gain and loss events constitutes a complex trade-off [3]: while the acquisition of genes enables the emergence of novel functions, it also may impose a metabolic burden on the organism, demanding energy and resources for the synthesis and maintenance of the acquired genes. Whether a gene is kept, is determined by the equilibrium between the fitness advantages within the specific environment and the metabolic cost associated. For instance, the presence of antibiotic resistance genes introduces a trade-off whereby a resistant strain will outcompete susceptible bacteria in the presence of antibiotics, but the metabolic burden in the absence of antibiotics will disadvantage the resistant strain [4–6]. These costs can be highly variable, with some resistance genes imposing little to no fitness cost or even providing a benefit [7].

Not only the presence of such mechanisms is fundamental for the evolution of a bacterial population, but the time component plays a key role in determining their effectiveness. The capacity of a bacterial community to swiftly gain or lose genes, termed genome plasticity, is an important factor contributing to its evolutionary success.

In the era of massive genomics data, investigating genome plasticity on both large and short time scales has become possible. To do so, however, *ad-hoc* tools must be designed. Over the years, several metrics have been proposed to assess genome plasticity and offer insights into the dynamics of genetic exchange among bacteria, such as Jaccard distance applied to gene content [8, 9]. Jaccard distance, a dissimilarity measure widely used in machine learning and computational genomics [10], can be applied to compare gene content across different sets of genomes using the following formula:

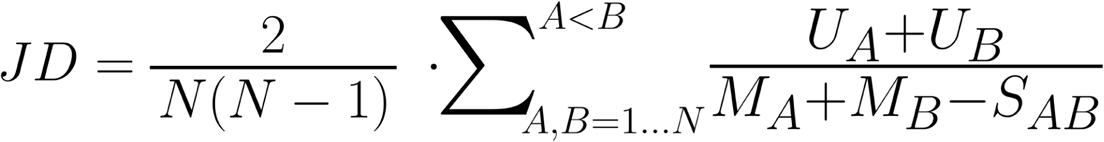

where U_A_ and U_B_ are the number of genes found only in genomes a and b respectively and M_A_ and M_B_ are the total number of genes found in a and b respectively, S_A,B_ are the shared genes between a and b, and N_p_ the total number of pairs considered.

In this context, Jaccard distance allows to compare the overall gene repertoire across different groups of genomes. Moreover, Jaccard distance applied to gene content resembles the genome fluidity formula [11]:

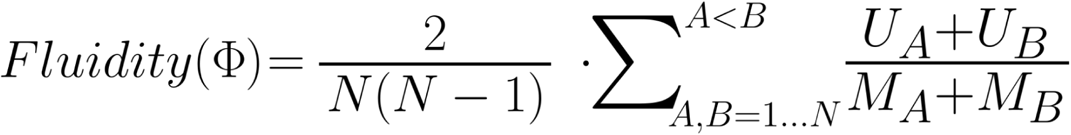

where U_A_ and U_B_ are the number of gene families found only in genomes A and B respectively, M_A_ and M_B_ are the total number of gene families found in a and b respectively, and N_p_ the total of genome pairs.

Although the difference is slight, when Jaccard distance is used to compare gene content between two genomes, it assigns lower importance to core genes than genome fluidity. This is achieved by preventing the double counting of shared gene families between the two genomes. Genome fluidity was proposed by the authors as an alternative to core and pan genome for measuring gene content diversity within a species or groups of closely related organisms [11].

Neither Jaccard distance nor genome fluidity take into account the time component, which as previously stated, is a key component in defining genome plasticity. As a consequence, by applying these two indexes to gene content it is not possible to distinguish between a scenario where a bacterial community gradually accumulates a large gene repertoire over an extended period and a second scenario where the population undergoes rapid gain or loss events. The latter is a distinctive feature of emerging high risk clones in many bacterial pathogen species, one commonly associated with increased virulence, enhanced transmissibility, and extensive antibiotic resistance [12–14]. Additionally, both indexes consider genes as single entities, whereas HGT events often involve regions with more than one gene. Consequently, even with a time correction, the number of gene gain or loss events may be overestimated.

In this study we aimed at providing a new metric to assess the rate at which bacteria exchange genes. We implemented a new method to compute the number of gene gain and loss events and introduced the evolutionary distance into the calculation. Then, we used the new index to investigate the dynamics of bacterial genome plasticity within three widespread, widely studied bacterial species: *Klebsiella pneumoniae*, *Staphylococcus aureus*, and *Escherichia coli*.

## Methods

### Datasets construction

A curated dataset of high quality genomes was collected from BV-BRC [40] (updated December 2022) independently for *K. pneumoniae*, *S. aureus*and *E. coli* using the makepdordb.py script of the P-DOR pipeline [41], which automatically filters for genome size and number of contigs. To further improve the quality of each dataset, we performed *in silico* Multilocus Sequence Typing (MLST), using schemes downloaded from PubMLST [42] in December 2022. For *E. coli* we chose the Achtman scheme. Since MLST is based on single-copy housekeeping genes, the absence, or the presence of more than one copy, of one of these genes is most likely owing to a poor quality genome assembly. Only genomes that could be assigned a Sequence Type (ST), using an *in-house* python script, were used in the next phase.

To reduce redundancy (e.g. to avoid almost identical genomes that were obtained to analyze outbreaks) from each dataset, dRep [43] was applied using the “dereplicate” function (-ms 10000 -pa 0.99 --SkipSecondary), also allowing to confirm the quality of the genome using its internal default pipeline.

A SNP alignment was obtained for the reduced datasets using P-DOR (-n 0) [41] and then used as input for fastBAPS [16] with default settings. After eliminating any fastBAPS-assessed clusters with less than 100 genomes, a random selection of 100 genomes was chosen from each cluster that was still present to ensure representativeness. We named each cluster with the prevalent ST; when a ST was split between clusters, we added a letter to the name.

### SNP distances

A SNP alignment was generated on the final datasets using P-DOR (-n 0) [41] with an internal complete genome as reference for each species (*E. coli*: CP043539; *S. aureus*: LT963437; *K. pneumoniae*: CP006648). Pairwise SNP distances were computed using the snp-dists tool (https://github.com/tseemann/snp-dists).

### Flux of Genes Segments

In order to compute FOGS, we followed the same approach of Brilli and colleagues (Brilli et al. 2013) by translating each genome into a graph. (1) PanTA [44] was used to classify all the proteins from each species dataset into orthology groups. (2) The gene neighborhood network of each genome was built using the information about coding genes location in the annotation file previously produced by prokka (Seemann 2014). Each gene was connected to the one downstream in the genome table only if their distance was less than 1000 bp and if they belonged to the same contig. We then performed the pairwise graphs (genomes) comparison as follows: (3) for each genome the unique genes are retrieved, (4) and compressed if consecutive, considering them as a single element. (5) The sum of the unique gene segments obtained corresponds to the number of gene gain or loss events. To this purpose, we used the connected_components function included in the SciPy python library (Virtanen et al. 2020). (6) The number of gene gain or loss events was then weighted on the SNP distance, computed as previously described. All the steps were performed using an *in-house* python script.

### Assessing FOGS robustness

To evaluate the effect of the number of genomes on FOGS, we randomly sampled N genomes from the *Klebsiella pneumoniae* BV-BRC dereplicated dataset, with N ranging from 10 to 1000. Between N=10 and N=100, selections were made in steps of 10, while larger datasets (N>100) were selected in steps of 100. For each N, genomes were sampled randomly, with repetitions performed 10 times to ensure robustness across replicates. This bootstrap approach allowed us to evaluate the stability of FOGS estimates across multiple iterations. Then, to evaluate the influence of genome fragmentation on FOGS, we focused on closed genomes (i.e., those assembled into a single contig) and artificially fragmented them. We selected two sets of 100 complete genomes, with no overlap. Fragmentation was simulated by splitting each genome into N pieces of varying sizes (from 10 to 1000) using an in-house Python script. This approach ensured that only the contiguity of the genome was altered, while the total number of genes and gene sequences remained unchanged. Finally, to investigate the effects of dataset inhomogeneity, we computed FOGS across curated subsets of 100 genomes. These subsets were selected to vary systematically in their levels of inhomogeneity by adjusting the proportion of complete fragmented genomes within each dataset.

### Resistance and Virulence genes

To assess the presence of resistance genes, we utilized Resfinder [45] (Florensa et al., 2022). Our criteria for determining gene presence required a minimum of 60% query coverage and 90% sequence identity. Similarly, we assessed the presence of virulence genes using the Virulence Finder Database (VFDB) [46]. Analyses on *K. pneumoniae* were repeated using Kleborate [47].

To ascertain the presence and variant of *fimH*, a dedicated BLASTn search was performed using the fimtyper database (bitbucket.org/genomicepidemiology/fimtyper_db) as reference (100% query coverage, 100% sequence identity).

### Statistical analysis

The mean distribution of the number of gene gain/loss events weighted by SNP distances (*X*_A,B_/d_A,B_) between fastBAPS clusters were tested using the Kruskal-Wallis test, followed by Dunn’s test for pairwise comparisons using Benjamini-Hochberg adjustment. The analyses were performed using R v4.1.3.

## Results and discussion

### A novel index: Flux of Genes Segments

To assess genome plasticity, here we introduce a novel index called Flux of Genes Segments (FOGS). This new index is designed to align with the aforementioned definition of genome plasticity, representing the rate of events of gene acquisition or loss. To achieve this, we developed a new method for calculating the number of such events and integrated the time component to express it as a rate.

To address the challenge of accounting for gene gain or loss events, we follow the most parsimonious hypothesis, thus when multiple neighboring genes are gained or lost, we count a single event rather than distinct ones. Our approach to calculate the number of events is presented in Figure 1. We represented each genome into a graph connecting consecutive genes, akin to the method employed in the computation of the Genome Organization Stability index [15]. When two genomes are compared, shared genes are excluded from the analysis, leaving only unique genes to be considered. This approach eliminates the influence of internal rearrangements, which would otherwise introduce noise into the calculation, as such rearrangements do not impact the number of gene gain or loss events relevant to the index. Exploiting tables of gene coordinates, if genes are consecutive they will be compressed and considered as single entities. By summing the total number of these unique gene stretches, we calculate the putative number of gene gain or loss events occurring between the two genomes (Figure 1). It is important to highlight that this approach cannot distinguish whether an event represents a gain in one genome or a loss in the other. Instead, the focus is on the net result of these changes. Detailed technical explanations of the methodology are provided in the Materials and Methods section.

**Figure 1.**
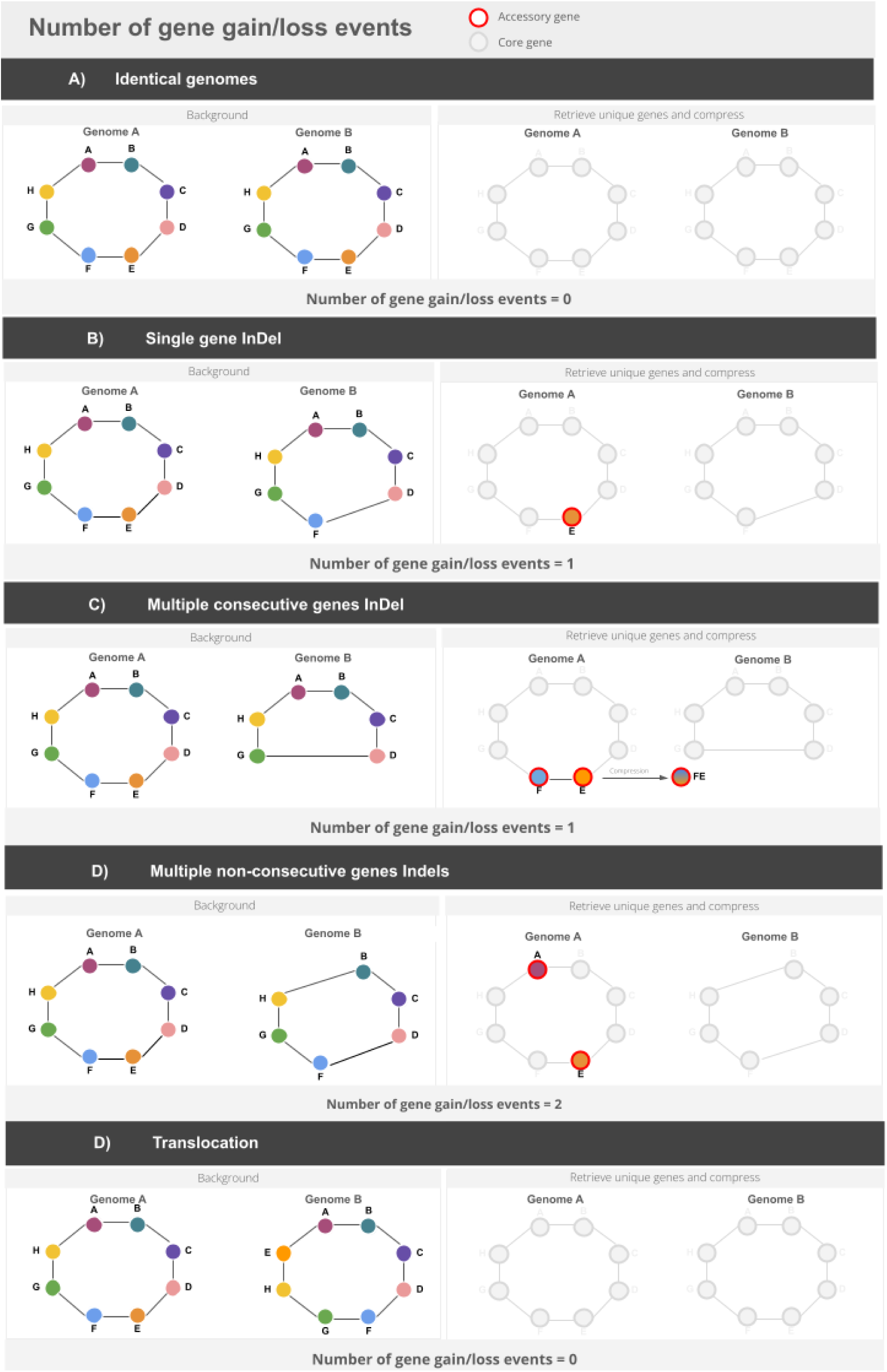
Strategy applied to compute the total number of genes gain/loss events considering different scenarios: A) Identical genomes, B) Single gene Indels, C) Multiple consecutive gene Indel, D) Multiple non-consecutive gene indels and E) Translocations.

Once the number of putative gene gain or loss events is determined for each pairwise comparison, we introduce temporal scaling by dividing the calculated value by the evolutionary distance between the genomes. For measuring evolutionary distance, we chose to use the core SNP distance. This decision was guided by our focus on applying the index to closely related strains, particularly in the context of intraspecific analyses. Core SNPs are particularly useful for evolutionary purposes since they preserve a higher discriminant power compared, for instance, to single copy orthologs (Bush et al. 2020). CoreSNPs may not be appropriate for studies involving entire species or comparisons across different species. In such cases, alternative approaches for measuring evolutionary distance, such as average nucleotide identity (ANI) or phylogenomic methods based on single-copy orthologs, may be more suitable.

Based on the above, for a group of *N* genomes FOGS is defined as:

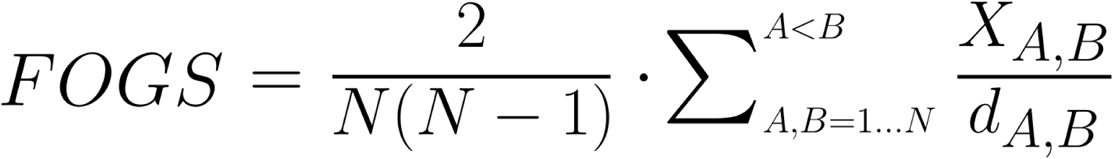

where *X* is the number of gene gain/loss events, *d* is the SNP distance between genome *A* and genome *B*, *N* is the number of genomes considered). The higher the value of FOGS, the higher is the genome plasticity.

### Assessing the robustness of FOGS

To ensure the reliability and applicability of the novel genomic plasticity index, we evaluated its sensitivity to key dataset characteristics, including the number of genomes and the level of genome fragmentation. Specifically, we examined how variations in these factors influence the accuracy and robustness of FOGS, identifying potential biases or limitations that could arise under different dataset conditions.

First, we tested datasets of varying sizes to determine the minimum sample requirements needed to produce consistent and meaningful results. The number of genomes in a dataset directly influences both the number of pairwise comparisons and the overall diversity captured in the analysis. To evaluate the impact of FOGS on dataset size, we examined multiple randomly selected datasets of varying sizes, ranging from small (N = 10) to larger populations (N = 1000). Our analysis demonstrated that the FOGS index exhibits stability when applied to medium and large datasets, with the index reaching a plateau at around N = 90 genomes. Increasing the number of genomes beyond this threshold does not substantially alter the outcome (Figure 2A). This finding highlights the critical importance of ensuring an adequate sample size to obtain robust and reproducible estimates of genomic plasticity, while also minimizing the influence of random variability. It also implies that smaller datasets may lead to less stable estimates, and including enough genomes in the analyses is highly recommended to avoid potential biases associated with under-sampling. We then investigated the impact of genome fragmentation, due to draft assemblies, on defining gene content and adjacency. Draft assemblies present a challenge for the FOGS index because they often consist of fragmented genomes represented by contigs, which are short, incomplete sequences of DNA that are not yet fully assembled into a complete genome. When working with contigs the order of genes is not always clear because the contigs are isolated segments that may not be placed in their correct relative positions in the genome. This lack of information about the gene order can lead to inaccuracies in calculating gene gain or loss events, as the adjacency between genes is a key component of how the index measures genomic plasticity. As a result, gene adjacency could be misrepresented, and the FOGS index might incorrectly assess gene gain or loss, as it is unable to account for the true order of the genes. To evaluate the impact of fragmentation, we simulated varying levels of fragmentation by dividing complete genomes into smaller contigs of different lengths. In Figure 2B, two distinct populations are observed, with FOGS values increasing as fragmentation levels rise. This indicates that, FOGS calculations for these artificially fragmented datasets overestimated gene gain/loss events when compared to the complete genomes, and the higher the fragmentation, the higher the overestimation (Figure 2B). This highlights the importance of preprocessing datasets to remove low-quality draft genomes whenever possible, to ensure more accurate and reliable FOGS calculations. In real-world applications, genomic datasets often present varying degrees of completeness and fragmentation, which can influence the reliability of FOGS. Our analysis showed that when fragmentation levels are relatively uniform within a dataset, the associated error follows a predictable linear relationship with the degree of fragmentation (Figure 2B). This consistency allows for straightforward comparisons between groups even with highly fragmented genomes. However, in datasets with high variability, particularly those dominated by fragmented genomes, we observed a significant increase in the variance of FOGS results (Figure 2C). To reduce these issues, we recommend avoiding comparing dataset with significantly different levels of fragmentation, since this will bias the index. With the current advancements in sequencing technology, the number of genomes and their quality are no longer the limiting factors they once were. Today, high-throughput sequencing methods and improved assembly techniques have made it possible to generate large numbers of high-quality genomes with reduced fragmentation. This significantly mitigates the impact of incomplete genomes and enhances the representativeness of datasets. As a result, large genomic populations can now be analyzed with greater reliability, enabling more accurate estimations of genomic plasticity, even in studies involving diverse genomic sets.

**Figure 2.**
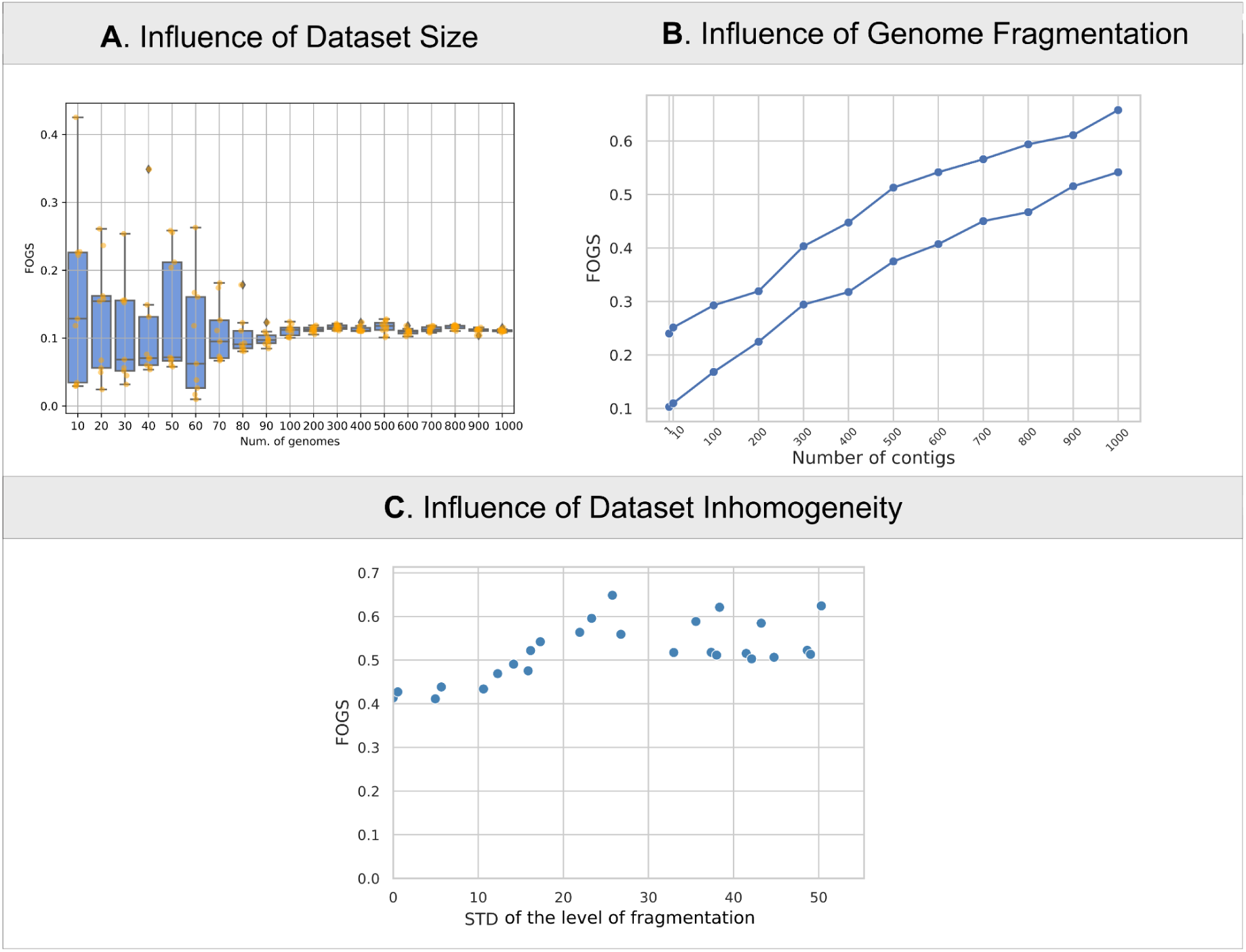
Robustness of FOGS to dataset characteristics. (A) The stability of the FOGS index (y-axis) was assessed by varying the number of genomes in the dataset (x-axis), ranging from 10 to 1000. For each dataset size, genomes were randomly selected 10 times, and the results are represented as boxplots. The orange points represent the actual FOGS values. (B) The influence of genome fragmentation on the accuracy of FOGS (y-axis) calculations was evaluated by selecting two random sets of 100 genomes and simulating different levels of fragmentation (x-axis). (C) The effect of dataset inhomogeneity, represented by the standard deviation of genome fragmentation (x-axis), on FOGS values (y-axis) was analyzed. This highlights the relationship between variability in genome completeness and FOGS reliability.

### Application of plasticity indexes to large genome datasets

We applied FOGS to investigate plasticity trends within three species of clinical interest. We choose as case studies *K. pneumoniae*, which is a well known nosocomial bacterium often associated with multidrug resistance, *S. aureus,* which is both nosocomial and community acquired, and *E. coli*, known for its wide repertoire of ecological niches. For each organism, we identified prominent taxonomic clusters using the fastBAPS algorithm [16] from a comprehensive collection of high-quality genomes accessible through the BV-BRC database [17]. Then, we purged nearly-identical genomes (i.e. outbreaks) and selected a representative subset of 100 genomes from each of the most prominent clusters (>100 genomes). For convenience of interpretation, we labeled each cluster with the most represented Multi-Locus Sequence Type (MLST or ST) among the genomes. See Table 1 for an overview of the results obtained at each stage of the filtering process (for technical details see the Materials and Methods section).

**Table 1.** Datasets building. * Number of genomes for which it was possible to obtain the ST. Δ Number of genomes after the dereplication step. & Clusters detected by fastBAPS comprised at least 100 genomes. # Number of genomes used for subsequent analysis.

| Species | HQ<br>genomes | MLST* | dRep<sup>Δ</sup> | N. of<br>clusters<sup>&</sup> | N. of genomes<sup>#</sup> |
| --- | --- | --- | --- | --- | --- |
| *Klebsiella pneumoniae* | 17887 | 17435 | 11196 | 19 | 1900 |
| *Staphylococcus aureus* | 18849 | 18188 | 11319 | 20 | 2000 |
| *Escherichia coli* | 42049 | 40805 (#1,<br>Atchman) | 22099 | 35 | 3500 |

We applied the newly created index FOGS to the three species datasets in order to investigate the presence of different plasticity patterns among the clones. Initially, we considered the presence of a relationship between the number of gene gain or loss events versus the number of SNP for each pairwise comparison (Supplementary Figure 1) within each species. Although a general relationship was observed—indicating that a higher SNP distance corresponds to a higher number of potential gene gain/loss events—there is a noticeable clustering at low SNP distances. This clustering reveals a broad spectrum of putative events in this range. Also, the relationship observed is non-linear.

In addition, we investigate a possible association between FOGS and virulence or resistance traits by comparing the index to the mean number of resistance and virulence genes for each cluster. As follows, we discuss the results obtained for each species in depth.

### Klebsiella pneumoniae

*K. pneumoniae* is one of the leading causes of hospital-acquired infections, especially among immunocompromised individuals, elderly patients, and those with chronic conditions. Within the hospital environment, this bacterium can persist on various surfaces, medical equipment, and in water sources. Thus, it poses a significant challenge in terms of antibiotic resistance, making it a persistent concern for healthcare settings [18]. The *K. pneumoniae* population shows a remarkable level of diversity, encompassing numerous distinct phylogenetic lineages or ‘clones’ [19]. These lineages can be defined as Multidrug Resistant (MDR), or hypervirulent based on the prevalence of determinants of resistance to antibiotics or virulence genes.

Wyres and colleagues analyzed the gene content diversity between MDR and hypervirulent clones and found a reduced genetic diversity in the hypervirulent ones. They achieved this result by calculating the pairwise Jaccard gene content distances among genomes belonging to a clone [8]. In our study, we focused on evaluating the rate at which genome content evolves. Most of the clusters in our dataset, including STs 681, 13, 218, 348, 231, 395, 65, and 29 displayed a generally low gene flow. However, a smaller subset of clusters, exclusively represented by STs 307, 101, and 15 exhibited higher values, suggesting greater plasticity (Figure 3).

**Figure 3.**
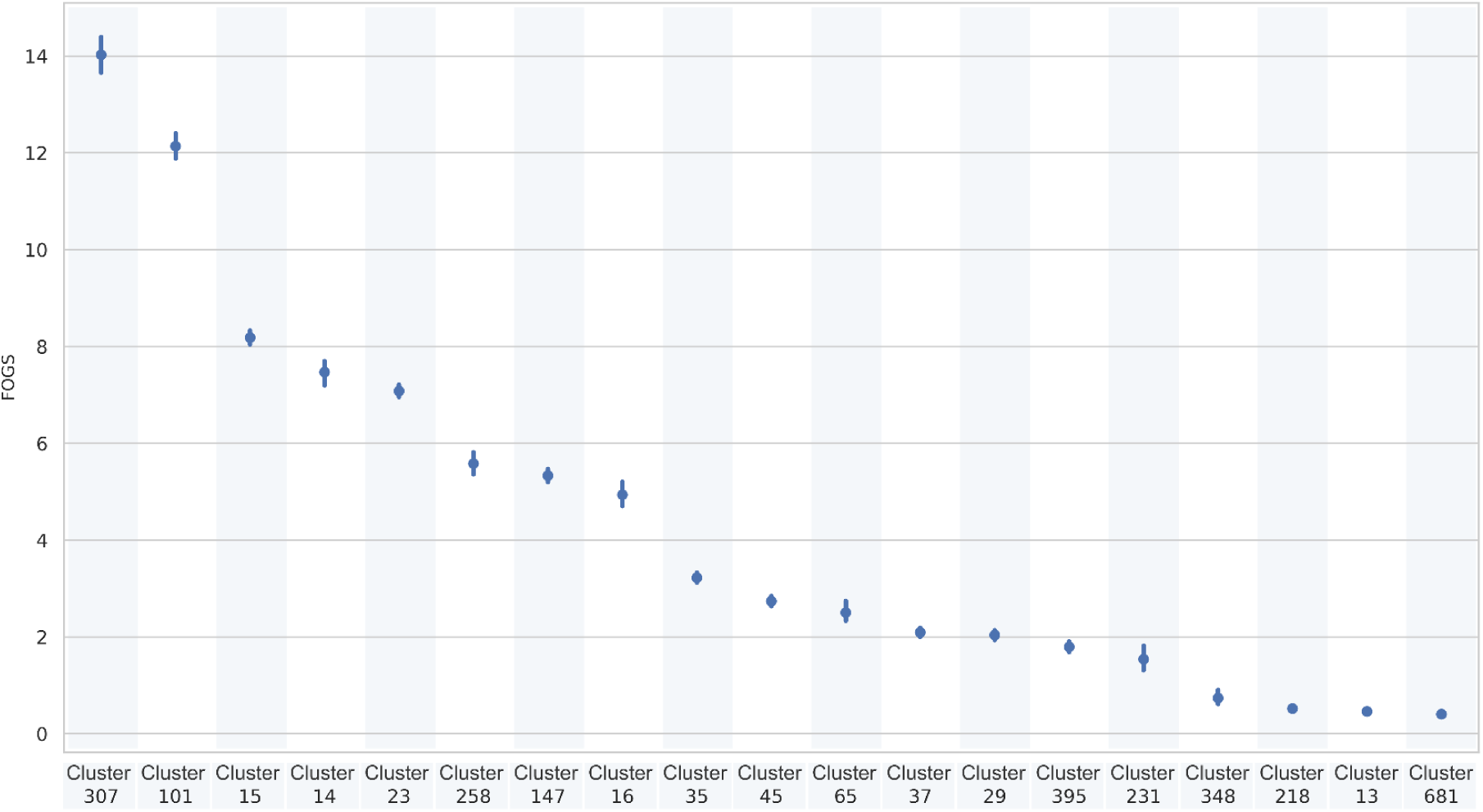
Pointplot FOGS within each cluster identified by fastBAPS [16] in *K. pneumoniae*. Vertical lines represent the confidence intervals.

Clusters at higher FOGS levels (307, 101, 15, 147) correspond to STs that are widely recognized globally as emerging high-risk clones, due to their potential to cause severe infections and their association with antimicrobial resistance (AMR), including pan resistance [12, 20–22]. Notably, ST307 has been gradually displacing the well known ST258 in multiple areas of the world (e.g.[12]). Readers should take into account that the cluster we named ‘258’ encompasses ST11, ST258 and ST512 which are part of the CG258. This lineage has been for a long time the most prevalent MDR-Carbapenemase producer group [20, 23].

Based on our analysis, cluster 307 exhibits a substantially higher genome plasticity compared to cluster 258 (p < 1E-10). This finding provides a possible explanation for the recent rise of ST307, which appears to be the most plastic. Moreover, considering the mean number of resistance genes per clone, ST307 stands out as one of the most resistant clusters in our dataset. A positive correlation between plasticity and antibiotic resistance can be observed (Supplementary Figure 2A, R^2^=0.44, p-value<0.05). This result is in accordance with the findings of Wyres and coworkers [8]. However, our results also suggest that no direct or inverse relationship is present between virulence and plasticity (R^2^=0.14, p-value>0.05) (Supplementary Figure 2B). To hypothesize an explanation to these observations, one should consider that virulence genes mostly imply a constant fitness advantage. So they are more likely to be maintained and transmitted vertically than resistance genes, which are only needed in the presence of the antimicrobial and repeatedly lost and regained (e.g.[24]). As a consequence, our results suggest that the *K. pneumoniae* strains with a higher plasticity are also more likely to be resistant to antimicrobials.

### Staphylococcus aureus

*S. aureus* is a versatile Gram-positive bacterium that colonizes the skin and mucous membranes of humans and animals. While it is a common member of the human microbiota, *S. aureus* can also cause a wide range of infections, from minor skin and soft tissue infections to life-threatening diseases such as bloodstream infections, pneumonia, and endocarditis [25]. As*K. pneumoniae*, *S. aureus* is known for its ability to acquire and maintain resistance to multiple antimicrobial agents, making it a significant public health concern worldwide [26]. Among resistant strains, methicillin-resistant *Staphylococcus aureus* (MRSA) is particularly challenging due to the limited availability of alternative treatment options [27]. Genetic factors such as *mec* genes are responsible for this resistance [28].

ST5 and ST8 are the two major STs in *S. aureus* and are commonly associated with various types of infections, including those acquired in healthcare settings as well as in the community [29, 30]. In our study, the fastBAPS algorithm successfully divided both ST5 and ST8 into distinct subgroups. We observed a noteworthy difference in plasticity within the subgroups of ST8, specifically the cluster 8A which exhibited significantly higher plasticity compared to the other subgroup, cluster 8B (p < 1E-10) (Figure 4). We hypothesized the presence of a highly successful and adaptable sub-strain within ST8, which exhibits the highest level of plasticity in our entire dataset. Subsequently, we investigated the presence of resistance and virulence determinants in these subgroups. Notably, cluster 8A was found to be enriched in specific resistance genes: *mecA* (responsible for methicillin resistance) was found in all the 8A genomes but only in the 60% of the 8B group (Supplementary Figure 3); *mph(C)* and *mrs(A)* confer resistance to macrolide antibiotics, such as erythromycin, azithromycin, and clarithromycin; *aph(3’)-III* is associated to the resistance to gentamicin, tobramycin, and amikacin; *ant(6)-Ia* provides resistance to aminoglycoside antibiotics, including kanamycin and neomycin. While the virulence pattern remained relatively stable between the two subgroups, we discovered a significant difference in the presence of CHIPS genes, which were found in the majority of genomes within cluster 8A. These genes play a crucial role as important virulence factors, helping *S. aureus* evade the innate immune defense systems [31]. These findings underscore the significance of cluster 8A as a particular plastic subgroup within the ST8. On the other hand, the two subgroups of ST5 did not show a significant difference in plasticity, resistance or virulence patterns.

**Figure 4.**
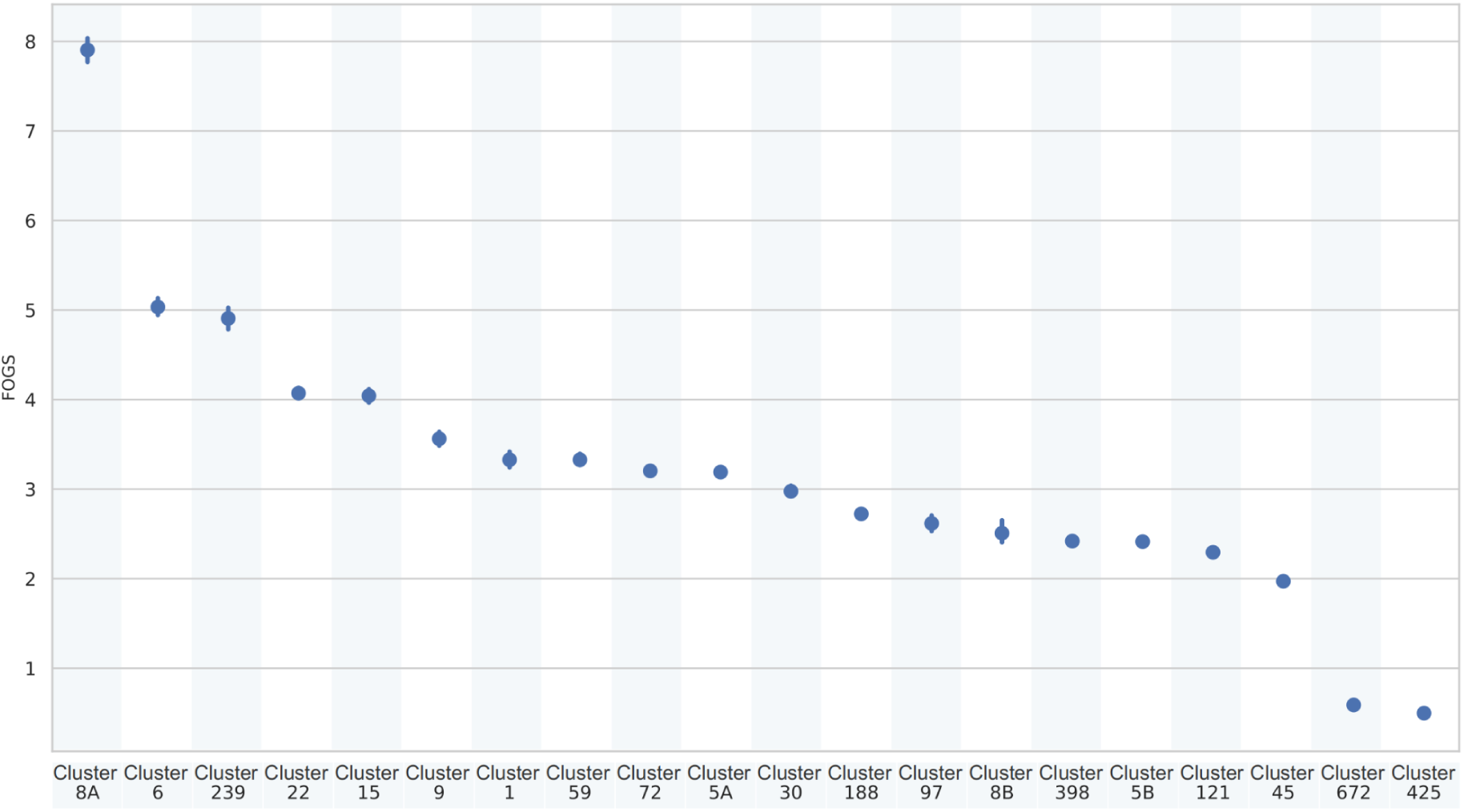
Pointplot FOGS within each cluster identified by fastBAPS [16] in *S. aureus*. Vertical lines represent the confidence intervals.

### Escherichia coli

*E. coli* is a widely studied Gram-negative bacterium that plays a crucial role in various ecological niches and has significant implications for human health. Previous studies have demonstrated the presence of extensive genetic variation within *E. coli*, enhanced by its ability to adapt to a range of diverse niches including hospitals, animal reservoirs, and natural ecosystems [32–34]. In the clinical setting, *E. coli*represents a significant public health concern due to its ability to cause a wide range of infections, ranging from urinary tract infections to more severe bloodstream infections.

ST131 is one of the most predominant and globally disseminated lineages associated with urinary tract infections (UTIs) and bloodstream infections. Frequently associated with multidrug resistance, including extended-spectrum beta-lactamases (ESBLs) and fluoroquinolone resistance, it can be considered the most successful MDR clone of all time [35, 36]. In our dataset, ST131 was split in two clusters, namely ‘131A’ and ‘131B’. Cluster ‘131B’ showed the highest FOGS value of the dataset, and significantly higher in respect to cluster ‘131A’ (p<0.01; Figure 5). ST131’s population structure has been previously investigated and three genetically distinct clades have been identified (A, B and C), each characterized by different fimbrial adhesin (*fimH*) gene variants [37]. Our analysis revealed that cluster 131A uniquely contains genomes encoding the *fimH*41 variant, which is exclusive to the globally recognized cluster A. In contrast, cluster 131B encompasses various *fimH* variants, including 30 and 40, thereby comprising the global clusters B and C. Within specific subclades of the global cluster C, a convergence of extensive resistance and virulence profiles has been observed [38]. Biggel and colleagues propose that this convergence may not be applicable to other *E. coli* lineages, such as ST73 and ST95. Despite being pandemic, these lineages exhibit low antibiotic resistance, potentially attributed to their gene acquisition capabilities [38]. Our findings align with this hypothesis, revealing a lower degree of genomic plasticity in ST73 and ST95.

**Figure 5.**
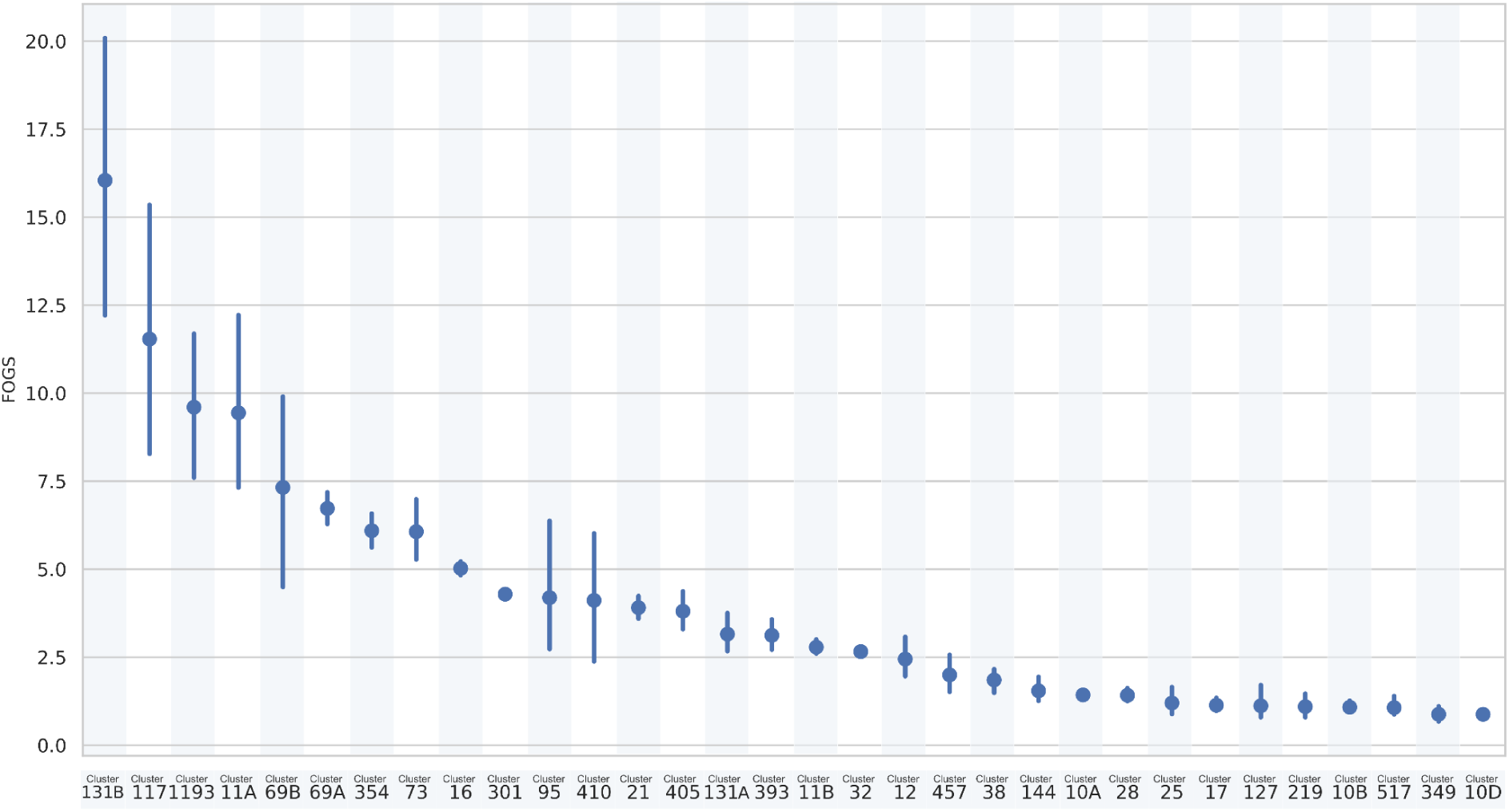
Pointplot FOGS within each cluster identified by fastBAPS [16] in *E. coli*. Vertical lines represent the confidence intervals.

Interestingly, cluster 1193 appears to be the second most plastic clone according to FOGS. *E. coli* ST1193 is currently emerging rapidly across the globe, mimicking the very successful ST131 [39].

## Conclusions

Gene gain and loss events represent a complex trade-off for prokaryotes, shaping their adaptive strategies in response to environmental challenges [3].

Here, we introduced a novel index, Flux of Genes Segments (FOGS), designed to quantify genome plasticity by assessing the rate of gene acquisition or loss events across multiple genomes. The integration of a temporal component in FOGS further emphasizes the importance of time in evaluating the effectiveness of adaptive mechanisms within bacterial populations. FOGS works with datasets containing either complete genomes or draft assemblies. The input data should ideally include high-quality gene annotations and clear gene coordinates, as FOGS relies on accurate identification of gene content and their adjacency within the genome. Additionally, datasets with varying levels of genome completeness and quality can be used, but it is recommended that caution be taken when dealing with heavily fragmented genomes, as they may introduce inaccuracies in the calculation of gene events.

We applied FOGS to analyze genome plasticity trends across three bacterial species: *Klebsiella pneumoniae*, *Staphylococcus aureus*, and *Escherichia coli*. Our results highlighted distinct plasticity patterns within specific sequence types and clusters, with notable connections to globally recognized high-risk clones. These findings suggest that FOGS can effectively capture variations in genome plasticity, offering insights into the evolution and adaptation of bacterial species, as well as their potential implications for public health and the emergence of antimicrobial resistance.

If validated further, FOGS could potentially allow for the early identification of strains of potential clinical concern (i.e. with a high rate of genomic plasticity), enabling more timely and targeted interventions to control the spread of antimicrobial resistance and prevent outbreaks of pathogenic strains.

Ultimately, the introduction of FOGS represents a powerful tool for the study of genomic plasticity, providing a quantitative measure that can be applied to a wide range of bacterial species. By incorporating the time factor, FOGS not only measures genetic events but also frames them within an evolutionary context, offering a more comprehensive view of how bacteria adapt and evolve over time.

## Author Statements

Conceptualization: S.G., G.B. and M.B. Methodology: S.G., G.B. and M.B. Formal analysis: G.B., S.G. and G.B.B. Software: G.B. and S.G. Writing – original draft: G.B., S.G. and D.S. Writing – review and editing: G.B., S.G., G.B.B., M.B., F.B. and D.S.

## Data availability

The scripts used to compute FOGS are available at https://github.com/MIDIfactory/Genome-Plasticity

