## Supplemental Figures 1,2,3 for "How to measure bacterial genome plasticity? A novel time-integrated index helps gather insights on pathogens"

### Supplementary Material

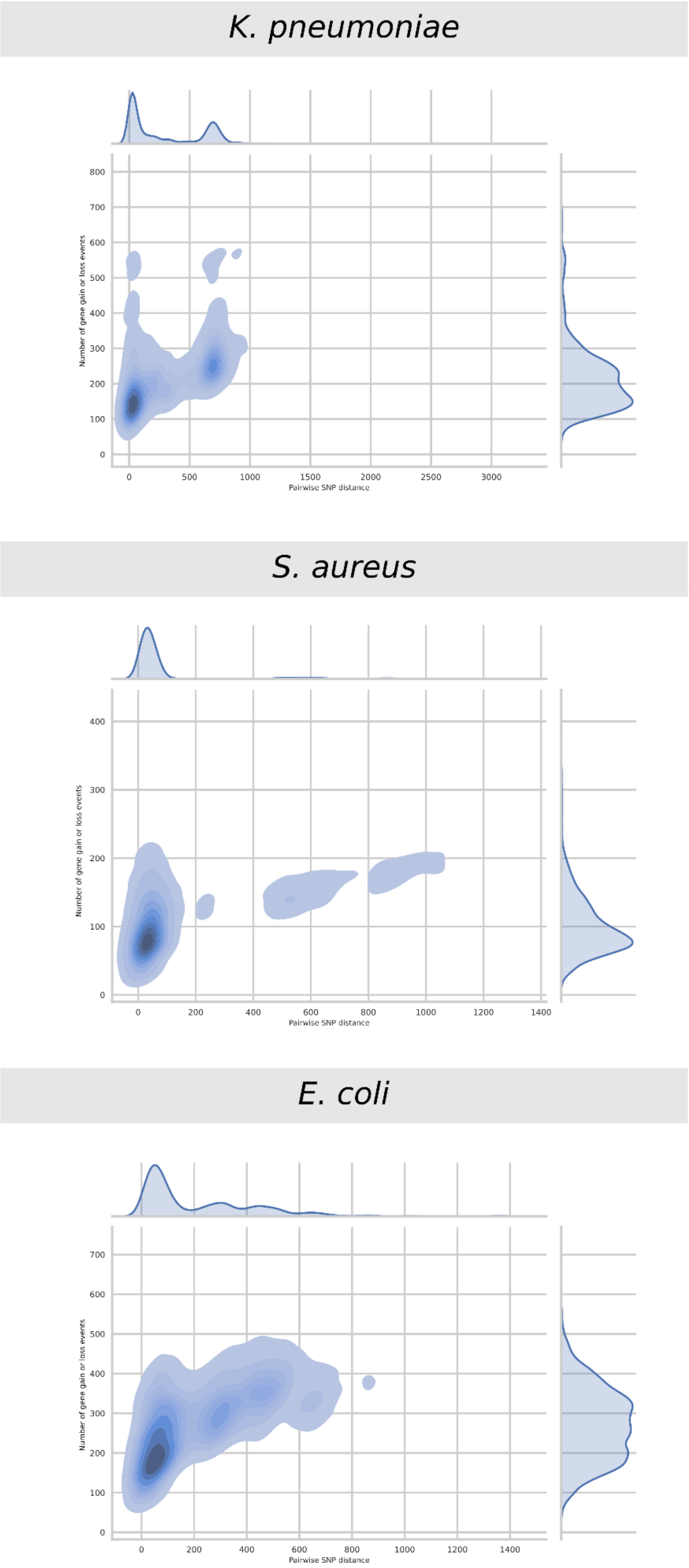


**Supplementary Figure 1**. Density plots of Number of gene gain or loss events against the SNP distance for all pairs within each fastBAPS cluster for *K. pneumoniae*, *S. aureus* and *E. coli*.

A

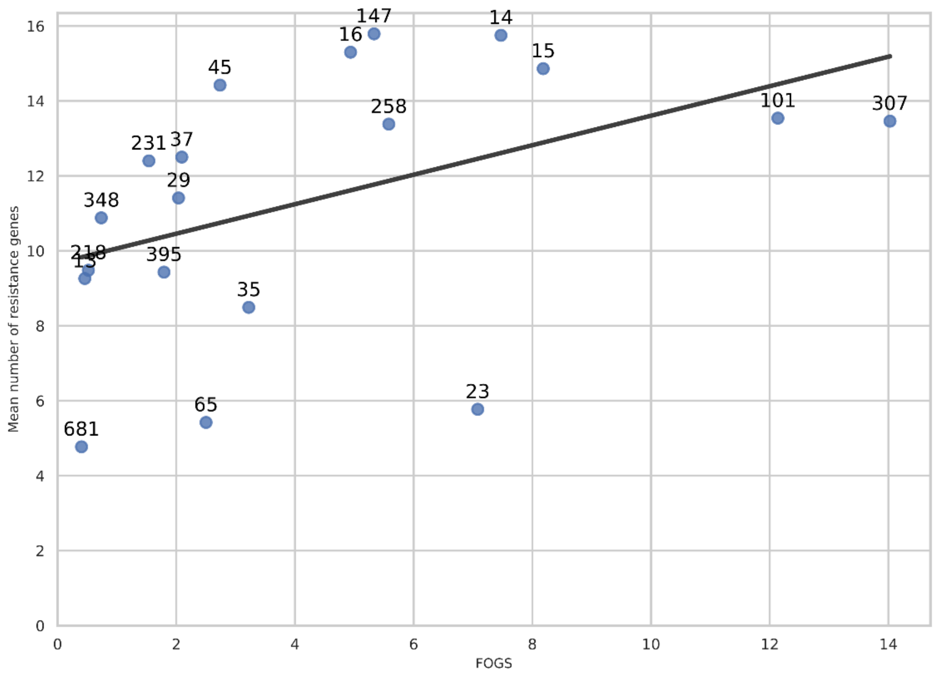


B

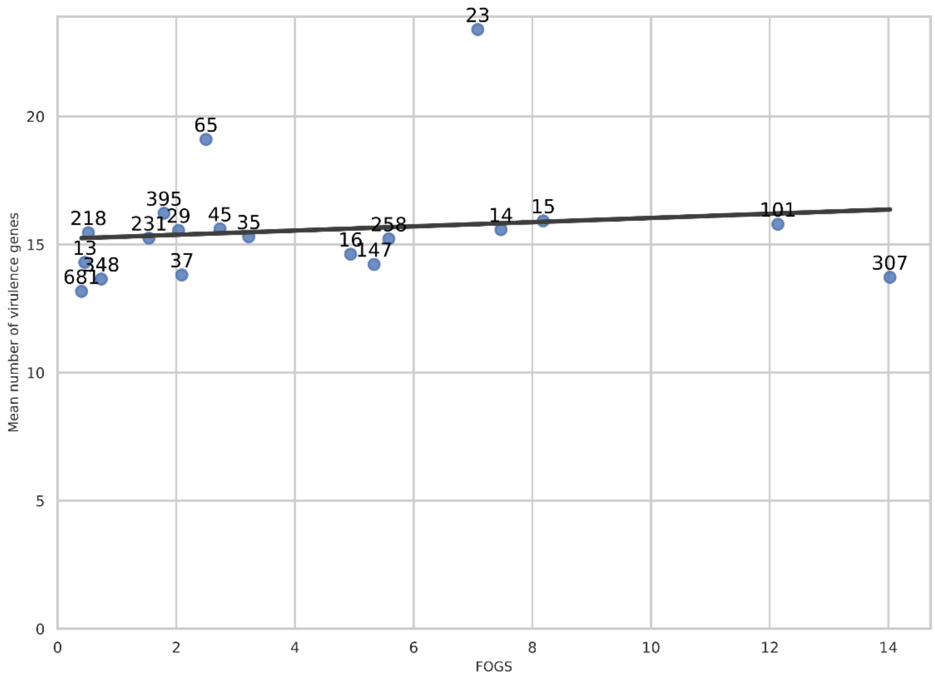


**Supplementary Figure 2**. A) Mean number of resistance genes against FOGS within each *K. pneumoniae* cluster. The regression line is presented in gray ($R^2$=0.44 , p-value<0.05). B) Mean number of virulence genes against FOGS within each *K. pneumoniae* cluster. The regression line is presented in gray ($R^2$=0.14 , p-value>0.05).

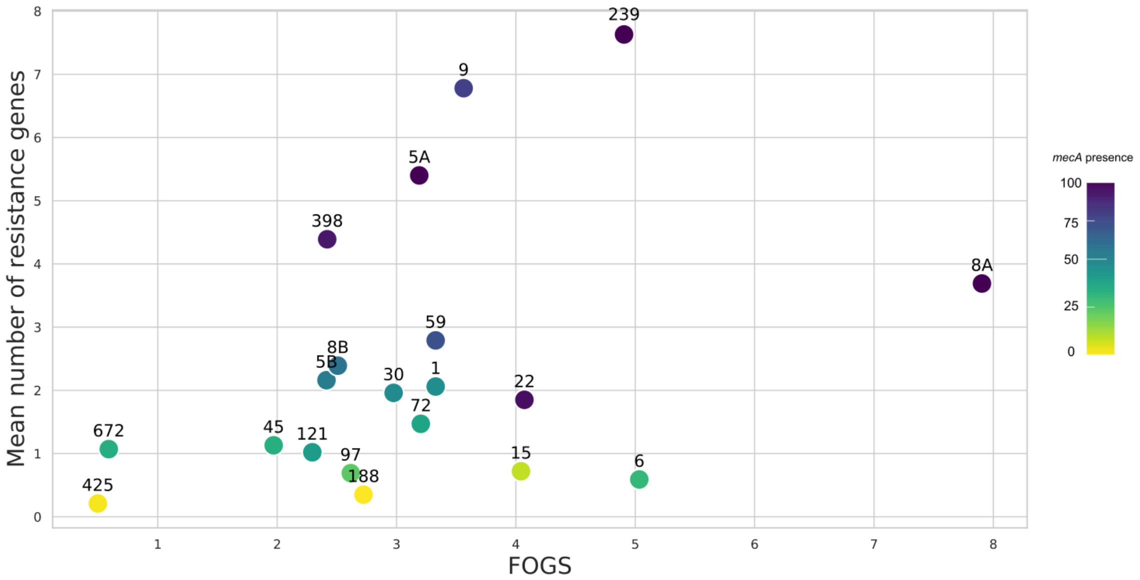


**Supplementary Figure 3**. Relationship between FOGS and the mean number of resistance genes in *Klebsiella pneumoniae*. The color of the point indicates the prevalence of the *mecA* within each cluster.
